# CytoLNCpred - A computational method for predicting cytoplasm associated long-coding RNAs in 15 cell-lines

**DOI:** 10.1101/2024.10.11.617765

**Authors:** Shubham Choudhury, Naman Kumar Mehta, Gajendra P. S. Raghava

## Abstract

The function of long non-coding RNA (lncRNA) is largely determined by its specific location within a cell. Previous methods have used noisy datasets, including mRNA transcripts in tools intended for lncRNAs, and excluded lncRNAs lacking significant differential localization between the cytoplasm and nucleus. In order to overcome these shortcomings, a method has been developed for predicting cytoplasm-associated lncRNAs in 15 human cell-lines, identifying which lncRNAs are more abundant in the cytoplasm compared to the nucleus. All models in this study were trained using five-fold cross validation and tested on an independent dataset. Initially, we developed machine and deep learning based models using traditional features like composition and correlation. Using composition and correlation based features, machine learning algorithms achieved an average AUC of 0.7049 and 0.7089, respectively for 15 cell-lines. Secondly, we developed machine based models developed using embedding features obtained from the large language model DNABERT-2. The average AUC for all the cell-lines achieved by this approach was 0.6604. Subsequently, we also fine-tuned DNABERT-2 on our training dataset and evaluated the fine-tuned DNABERT-2 model on the independent dataset. The fine-tuned DNABERT-2 model achieved an average AUC of 0.6336. Correlation-based features combined with ML algorithms outperform LLM-based models, in the case of predicting differential lncRNA localization. These cell-line specific models as well as web-based service are available to the public from our web server (https://webs.iiitd.edu.in/raghava/cytolncpred/) .

**HIGHLIGHTS:**

- Prediction of cytoplasm-associated lncRNAs in 15 human cell lines
- Machine learning using composition and correlation features
- DNABERT-2 embeddings for lncRNA localization prediction
- Correlation-based models outperform LLM-based models
- Web server and models available for public use

**AUTHOR’S BIOGRAPHY:**

1. Shubham Choudhury is currently working as Ph.D. in Computational Biology from Department of Computational Biology, Indraprastha Institute of Information Technology, New Delhi, India.
2. Naman Kumar Mehta is currently working as Ph.D. in Computational Biology from Department of Computational Biology, Indraprastha Institute of Information Technology, New Delhi, India.
3. Gajendra P. S. Raghava is currently working as Professor and Head of Department of Computational Biology, Indraprastha Institute of Information Technology, New Delhi, India

## INTRODUCTION

The rapidly expanding field of non-coding RNAs has revolutionized our understanding of gene regulation and cell biology. Among the diverse classes of non-coding RNAs, long non-coding RNAs (lncRNAs) have attracted significant attention due to their ability to regulate gene expression at various levels. Initially dismissed as transcriptional noise, lncRNAs have emerged as critical players in cellular processes, including development, differentiation, and disease progression [1]. To fully comprehend the functional roles of lncRNAs, it is imperative to investigate their subcellular localization. lncRNAs have distinct functions in the nucleus and cytoplasm, influencing transcriptional and posttranscriptional processes. In the nucleus, lncRNAs regulate gene expression and chromatin organization, while in the cytoplasm, they participate in signal transduction and translation. Some lncRNAs exhibit dual localization and functional diversification, reflecting their adaptability to different subcellular environments [2–4].

In recent years, extensive research efforts have been focused on deciphering the subcellular localization of lncRNAs. Various experimental approaches, such as fluorescence in situ hybridization (FISH) [5], RNA sequencing (RNA-seq) [6], and fractionation techniques [4], have been employed to identify the subcellular localization patterns of lncRNAs. These studies have revealed that lncRNAs can be localized in different cellular compartments, including the nucleus, cytoplasm, nucleolus, and specific subcellular structures. The subcellular localization of lncRNAs is often associated with their biological functions. For instance, nuclear-localized lncRNAs are frequently involved in transcriptional regulation, chromatin remodeling, and epigenetic modifications. Cytoplasmic lncRNAs, on the other hand, can interact with proteins or act as competitive endogenous RNAs (ceRNAs) to regulate gene expression post-transcriptionally [7]. However, most of these methods are expensive to perform and require highly specialized instrumentation.

Advancements in computational methods and machine learning approaches have further facilitated the prediction of lncRNA subcellular localization. These methods leverage various features, such as sequence composition, secondary structure, and evolutionary conservation, to predict the subcellular localization of lncRNAs with high accuracy. Several computational methods have been proposed for predicting lncRNA subcellular localization. Sequence-based methods rely on the nucleotide composition of the lncRNA. They utilize features such as k-mer frequency, nucleotide composition, and sequence motifs. However, these methods are trained on datasets that are not unique to humans, and they do not account for the variation in the subcellular localization of lncRNA in different cells.

Cell-line specific subcellular localization gains prominence due to the variability (in terms of subcellular localization) that lncRNAs exhibit within different cell-lines. This was reported by Lin et al. in lncLocator 2.0, where it was observed that a single lncRNA had different localization in different cell-lines [8]. We observed a similar trend in our dataset, where some lncRNAs were found to be localized in the nucleus for some cell-lines but were localizing to the cytoplasm in some other cell-lines. This pattern can be seen clearly in Figure 1.

**Figure 1.**
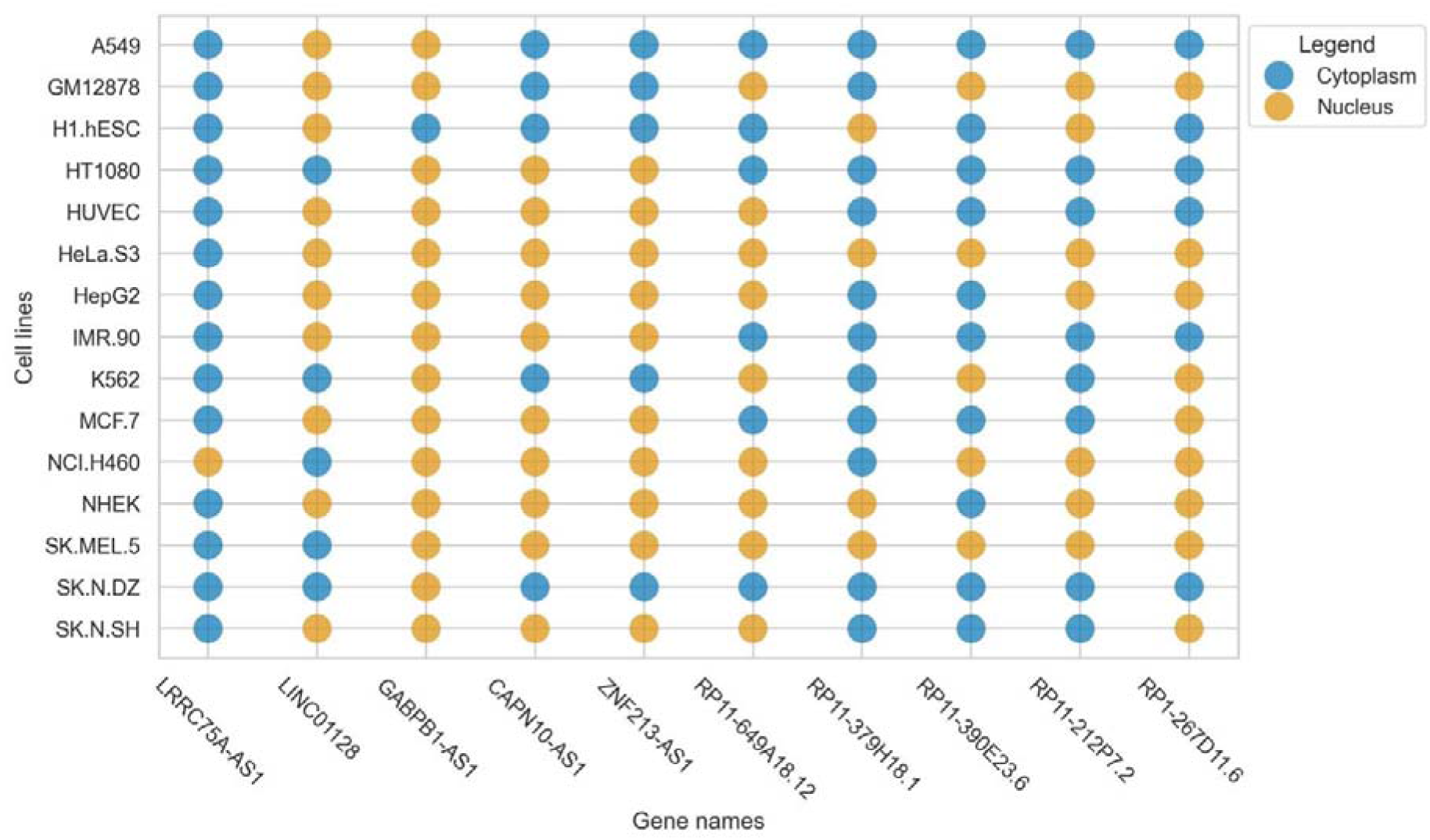
Bubble plot indicating the variability of localization of a single lncRNA across multiple cell-lines

**Figure 2.**
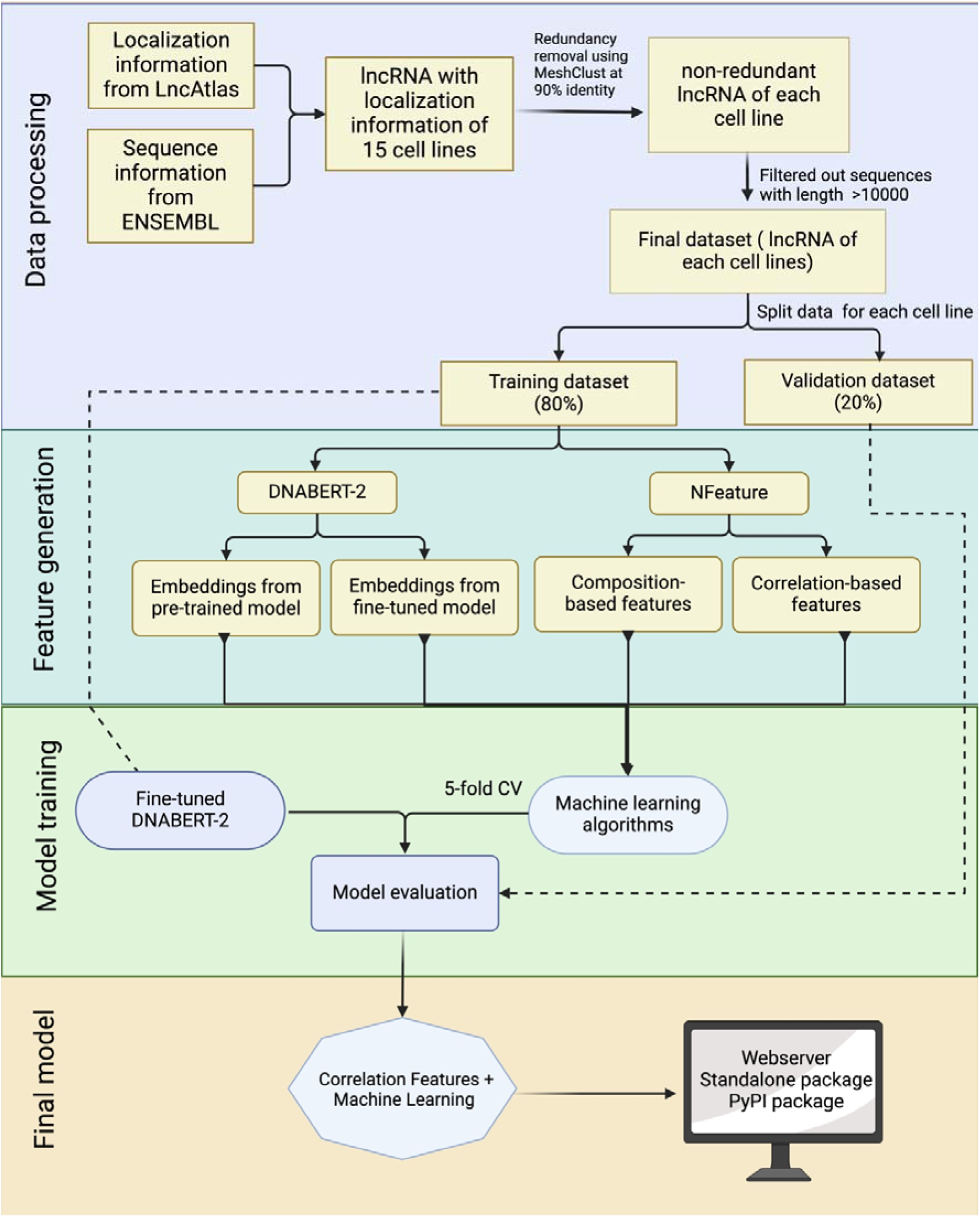
Overall architecture of CytoLNCpred

lncLocator 2.0 is a cell-line-specific subcellular localization predictor that employs an interpretable deep-learning approach [8]. TACOS, also a cell-line-specific subcellular localization predictor, uses tree-based algorithms along with various sequence compositional and physicochemical features [9]. Among all the existing computational methods, only lncLocator 2.0 and TACOS are designed to predict subcellular localization specific to different cell-lines. The primary issue with these methods is that the datasets used to develop these methods have not been properly filtered. Specifically, these methods have included mRNA sequences in their datasets, which can lead to inaccurate predictions. Additionally, the datasets have eliminated lncRNAs with an absolute fold-change less than 2, which can result in the failure to predict the subcellular location of lncRNAs with borderline concentration differences between locations.

To address the limitations of existing methods in a comprehensive manner, we have developed CytoLNCpred. In this study, we aimed to enhance the prediction accuracy compared to current tools, which have significant room for improvement. Furthermore, we have cleaned the dataset and adhered to industry standards to validate the performance of our method. In CytoLNCpred, a machine learning model trained using correlation-based features demonstrated significantly better performance on the validation dataset compared to existing tools.

## MATERIALS AND METHODS

To aid in the development of a prediction model for lncRNA subcellular localization, we’ve designed a workflow diagram, depicted in Figure 1. The comprehensive details of each phase in this workflow are outlined in the subsequent sections.

### Dataset creation

In this study, we have selected lncAtlas for acquiring cell-line specific subcellular localization information. lncAtlas is a comprehensive resource of lncRNA localization in human cells based on RNA-sequencing data sets [10]. lncAtlas contains a wide array of information, including Cytoplasm to Nucleus Relative Concentration Index (CNRCI), which we have utilized in our method. CNRCI is defined as the log2-transformed ratio of RPKM (Reads Per Kilobase per Million mapped reads) in two samples, in this case - the cytoplasm and nucleus. It is calculated as follows

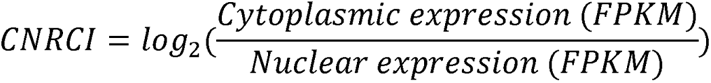

Sequence information for the lncRNAs was obtained from ENSEMBL database and lncRNAs with no sequence were dropped. In order to modify the dataset for a classification problem, we assigned sequences having CNRCI value greater than 0 as Cytoplasm and those having CNRCI value less than 0 were assigned as Nucleus. Redundancy was removed using MeshClust [11], using a sequence similarity of 90%.

Further, we used sequences up to the length of 10000 nucleotides only, as the longer lncRNA were misleading for the machine learning models and computationally very expensive when large language models were involved. The summary of the dataset used for each cell line is provided in Table 1.

**Table 1.** Detailed summary of the dataset used in the study, including the total number of samples for each cell-line in the source database and the final non-redundant dataset.

| Cell-lines | Original | Non-Redundant Dataset | Filtering out sequences > 10000 length | Training Dataset | Validation Dataset |
| --- | --- | --- | --- | --- | --- |
| A549 | 1826 | 1821 | 1131 | 904 | 227 |
| H1.hESC | 4194 | 4178 | 2552 | 2041 | 511 |
| HeLa.S3 | 1142 | 1141 | 703 | 562 | 141 |
| HepG2 | 1703 | 1699 | 1029 | 823 | 206 |
| HT1080 | 1183 | 1180 | 690 | 552 | 138 |
| HUVEC | 1870 | 1859 | 1137 | 909 | 228 |
| MCF.7 | 2714 | 2703 | 1702 | 1361 | 341 |
| **NCLH460** | 772 | 769 | 460 | 368 | 92 |
| **NHEK** | 1383 | 1378 | 755 | 604 | 151 |
| **SK.MEL.5** | 694 | 691 | 380 | 304 | 76 |
| **SK.N.DZ** | 762 | 759 | 422 | 337 | 85 |
| **SK.N.SH** | 2086 | 2077 | 1211 | 968 | 243 |
| **GM12878** | 2136 | 2128 | 1286 | 1028 | 258 |
| **K562** | 1197 | 1191 | 729 | 583 | 146 |
| **IMR.90** | 497 | 496 | 314 | 251 | 63 |

### Feature generation - Composition and Correlation-based

For facilitating the training of machine learning (ML) models, we generated a large variety of features using different approaches. These features convert nucleotide sequences We used the in-house tool Nfeature [12] for generating multiple composition and correlation features.

Composition-based Nucleotide composition-based features refer to quantitative representations of sequences that can be derived from the proportions and arrangements of nucleotides within these sequences. In this study, we have computed nucleic acid composition, distance distribution of nucleotides (DDN), nucleotide repeat index (NRI), pseudo composition and entropy of a sequence. The details for each of the features are provided in Table 2.

**Table 2.** Overview of the composition-based features generated using Nfeature.

| Feature name | Type of descriptor | No. of descriptors |
| --- | --- | --- |
| Nucleic acid Composition | Di-Nucleotide | 16 |
| Reverse complement K-Mer composition | Di-Nucleotide | 10 |
| Nucleotide Repeat Index | Mono-Nucleotide | 4 |
| Entropy | Sequence-level | 1 |
| Entropy | Nucleotide-level | 4 |
| Distance Distribution | Mono-Nucleotide | 4 |
| Pseudo Composition | Pseudo Di-nucleotide | 19 |
| Pseudo Composition | Pseudo Tri-nucleotide | 65 |
|  | **Total number of descriptors** | 123 |

### Correlation-based features

In this study, using Nfeature, we quantitatively assess the interdependent characteristics inherent in nucleotide sequences through the computation of correlation-based metrics. Correlation refers to the degree of relationship between distinct properties or features; an autocorrelation denotes the association of a feature with itself, whereas a cross-correlation indicates a linkage between two separate features. By employing these correlation-based descriptors, we effectively normalize the variable-length nucleotide sequences into uniform-length vectors, rendering them amenable to analysis via machine learning algorithms. These specific descriptors facilitate the identification and extraction of significant features predicated upon the nucleotide properties distributed throughout the sequence, enabling a more robust understanding of genetic information. A brief description of the features has been provided in Table 3.

**Table 3.**
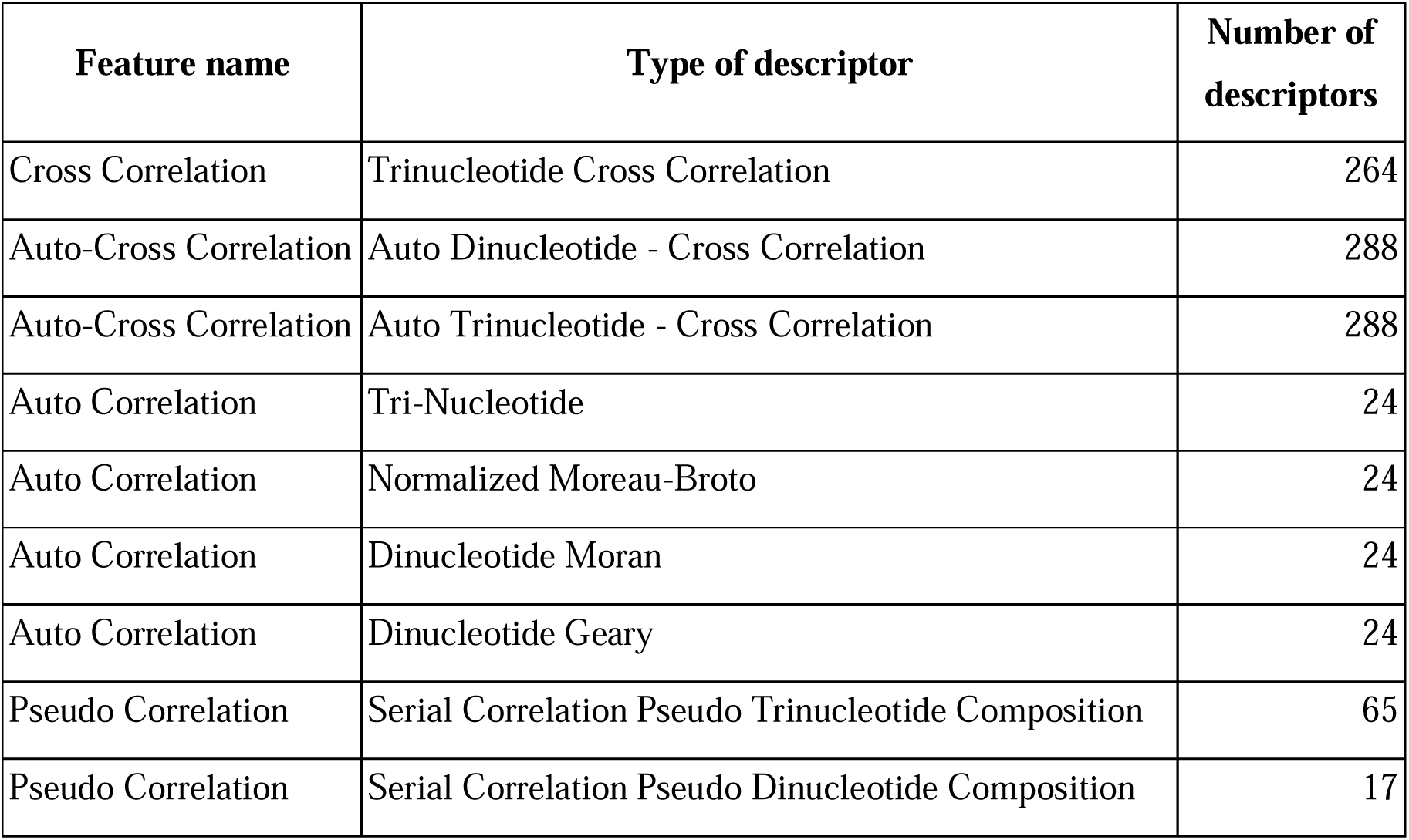
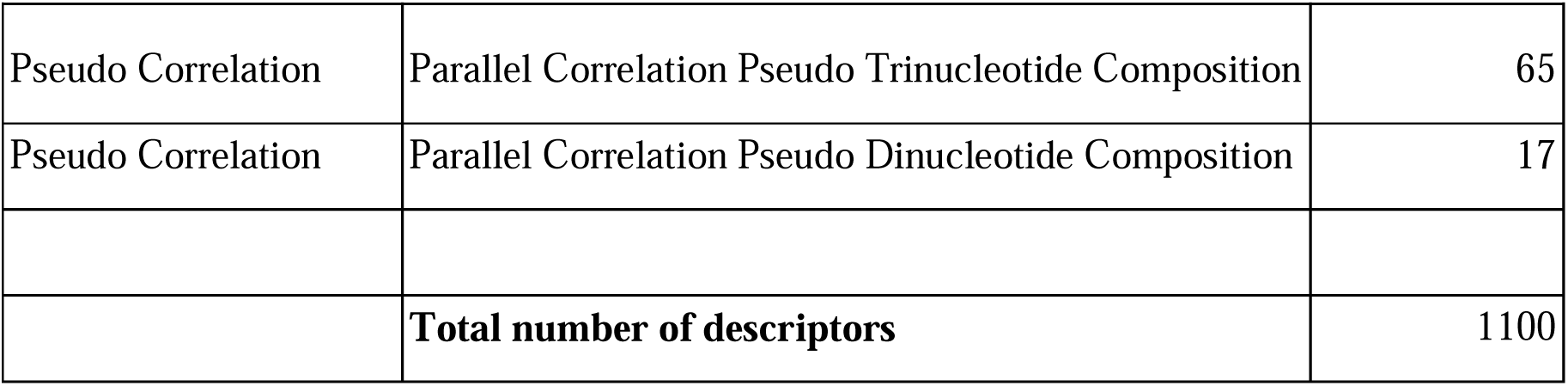
Overview of the correlation-based features generated using Nfeature.

| Feature name | Type of descriptor | Number of descriptors |
| --- | --- | --- |
| Cross Correlation | Trinucleotide Cross Correlation | 264 |
| Auto-Cross Correlation | Auto Dinucleotide - Cross Correlation | 288 |
| Auto-Cross Correlation | Auto Trinucleotide - Cross Correlation | 288 |
| Auto Correlation | Tri-Nucleotide | 24 |
| Auto Correlation | Normalized Moreau-Broto | 24 |
| Auto Correlation | Dinucleotide Moran | 24 |
| Auto Correlation | Dinucleotide Geary | 24 |
| Pseudo Correlation | Serial Correlation Pseudo Trinucleotide Composition | 65 |
| Pseudo Correlation | Serial Correlation Pseudo Dinucleotide Composition | 17 |
| Pseudo Correlation | Parallel Correlation Pseudo Trinucleotide Composition | 65 |
| Pseudo Correlation | Parallel Correlation Pseudo Dinucleotide Composition | 17 |
|  | **Total number of descriptors** | 1100 |

The total number of descriptors generated by using both composition and correlation-based features is 1223.

### Embedding using DNABERT-2

DNABERT-2 is an adaptation of BERT (Bidirectional Encoder Representations from Transformers) designed specifically for DNA sequence analysis [13]. DNABERT-2 generates embeddings for DNA sequences that encapsulate not just the individual bases, but also their biological significance in terms of structure, function, and interactions. Moreover, the key advantage of DNABERT-2 embeddings lies in their ability to capture the complex dependencies within DNA sequences. We have made use of both aspects of DNABERT-2 - the pre-trained model to make predictions and the embeddings from the model to be used as features for downstream tasks. The pre-trained model was trained on the training dataset using the default parameters mentioned in their GitHub repository. Moreover, we have also generated embeddings from both the pre-trained as well as the fine-tuned models in order to use them as features for machine learning algorithms. The number of embeddings generated for each lncRNA using DNABERT-2 was 768.

### Five-fold cross validation

In order to estimate the performance of models while training, we have deployed five-fold cross validation. In this method, the training dataset is split into five folds in a stratified manner and training is actually done over four folds and one fold is dedicated for validation. This process is iteratively performed for five times, by changing the fold that is used for validation and the rest of the folds being used for training. This generates an unbiased set of five performance metrics and the performance of the model is reported as the mean of these five sets.

### Model Development

In this study, three different approaches were followed for model development. The first approach involves the fine tuning of the DNABERT-2 using our training dataset and subsequently using the fine-tuned model to make predictions on the validation dataset. This method initially fine-tunes both the tokenizer and the pre-trained model according to our training dataset, and generates a fine-tuned tokenizer, and model. The final fine-tuned model takes lncRNA sequences and generates the prediction using the tokenizer and model. In the next approach, we have implemented a hybrid approach, combining the components of large language models and machine learning algorithms. Instead of features, we have generated embeddings from a pre-trained as well as fine-tuned DNABERT-2 model. Embeddings from the DNABERT-2 model were then used to train machine learning models and subsequently evaluated them.

### Model evaluation metrics

The binary classification performance of our fine-tuned model was evaluated using the following metrics: Sensitivity (SENS), Specificity (SPEC), Precision (PREC), Accuracy (ACC), Matthew’s Correlation Coefficient (MCC), F1-Score (F1) and Area Under the Receiver Operator Characteristic curve (AUC). The aforementioned metrics were calculated using the four different types of prediction outcomes: true positive (TP), false positive (FP), true negative (TN), and false negative (FN):

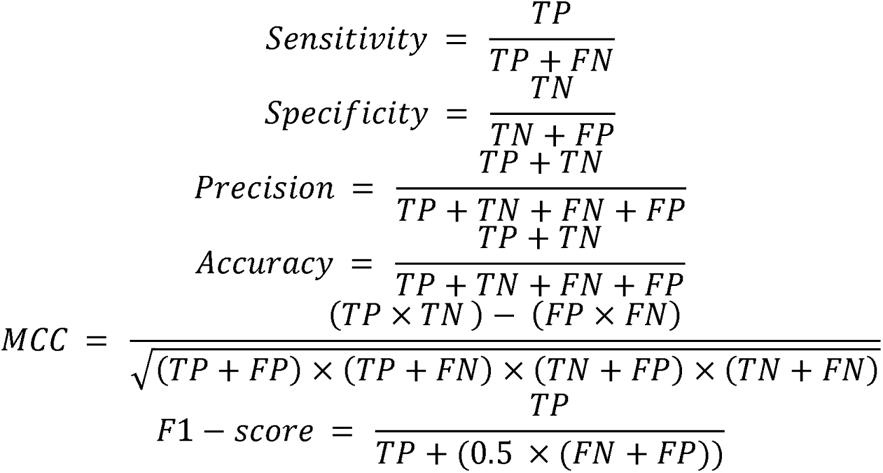

The evaluation of our binary classification model using various metrics provides critical insights into its performance. Sensitivity (SENS) measures the model’s ability to identify positive instances, while Specificity (SPEC) assesses its accuracy in recognizing negative instances. Precision (PREC) reflects the accuracy of positive predictions, and Accuracy (ACC) offers an overall measure of correctness, though it may be misleading in imbalanced datasets. Matthew’s Correlation Coefficient (MCC) provides a balanced view by considering all prediction outcomes, with values close to 1 indicating strong predictive capability. The F1-Score (F1) combines Precision and Sensitivity into a single metric, ideal for balancing the trade-off between false positives and negatives. Finally, the Area Under the Curve (AUC) evaluates the model’s ability to distinguish between classes across different thresholds, with higher values indicating better performance. Together, these metrics enable a comprehensive evaluation of the model, guiding necessary improvements and refinements.

## RESULTS

In this study, an attempt was made to design a model that will be able to classify the subcellular location of lncRNA into cytoplasm or nucleus. To achieve this, we tried out multiple approaches.

### Model based on composition and correlation features

Composition and correlation features generated from Nfeature were used to train multiple ML models. We have computed the performance of nine composition-based features and thirteen correlation-based features. We implemented all the combinations of feature and ML model to identify which feature-ML model combination performs the best. The composition features combined with classical ML methods were able to achieve an average AUC of 0.7049 and a MCC of 0.1965, across the 15 cell-lines. Similarly, with correlation-based features and ML methods, the best performance achieved was an average AUC of 0.7089 and a MCC of 0.2133. Performance of the best performing model using both composition and correlation-based features are provided for all the 15 cell-lines in Table 4 and 5 respectively.

**Table 4.**
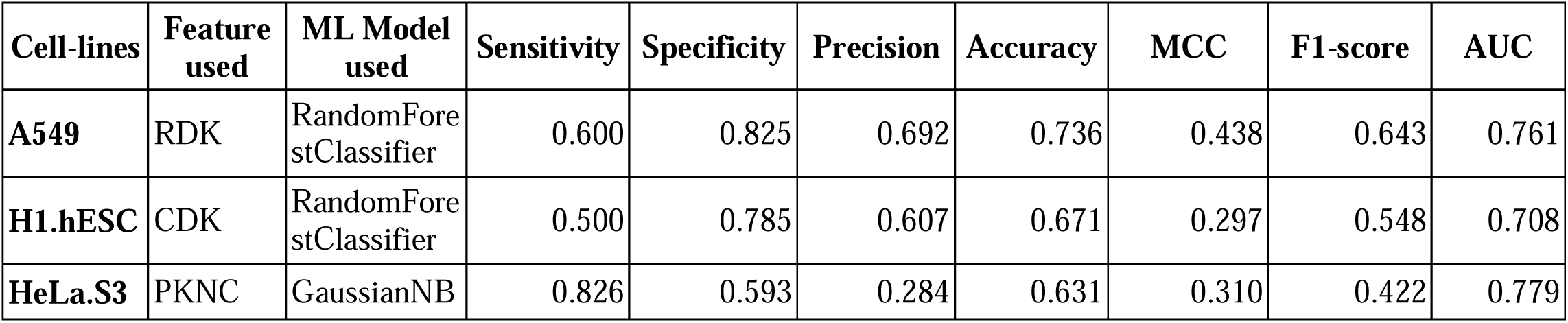
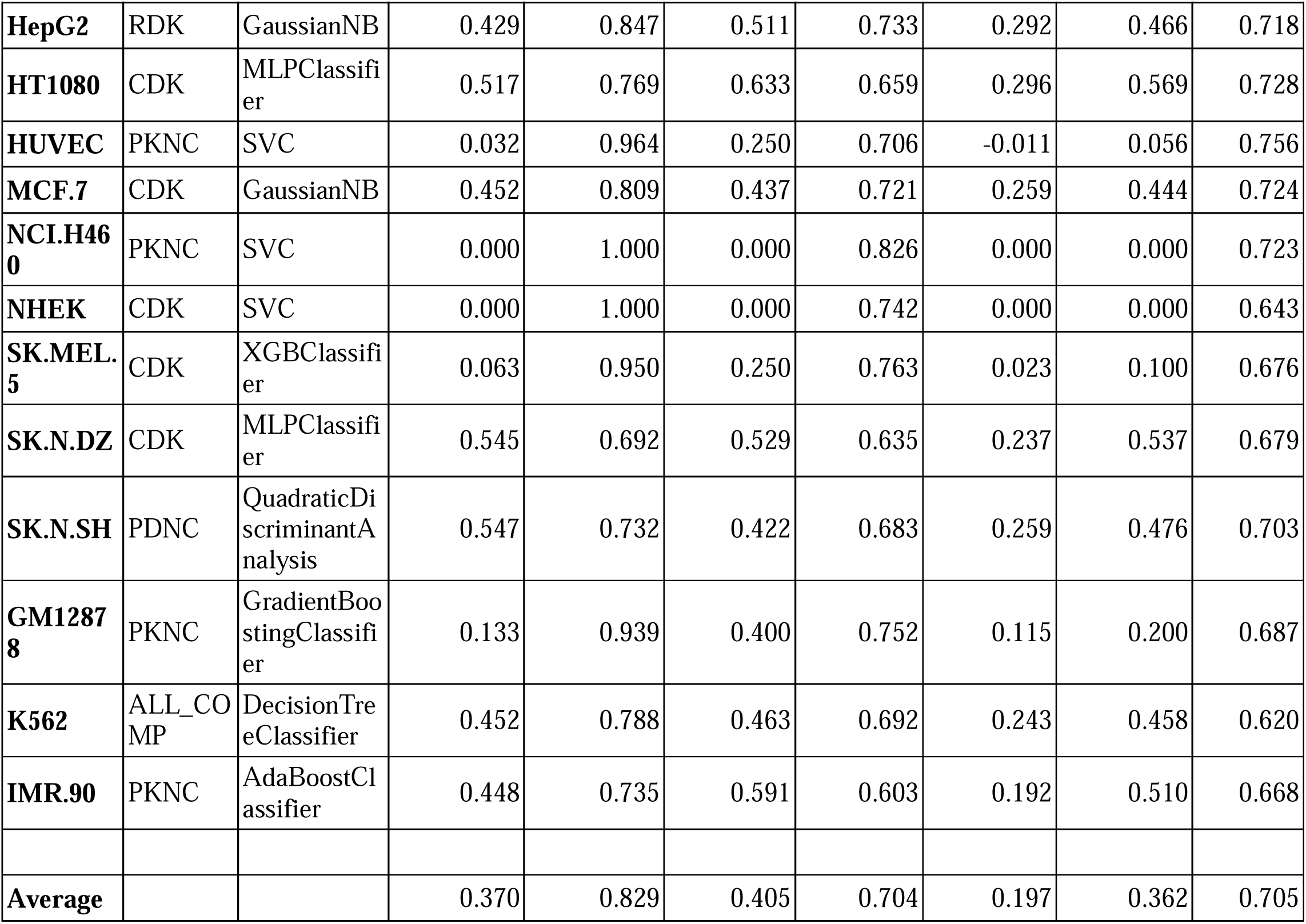
Performance of ML models on composition features.

| Cell-lines | Feature used | ML Model used | Sensitivity | Specificity | Precision | Accuracy | MCC | F1-score | AUC |
| --- | --- | --- | --- | --- | --- | --- | --- | --- | --- |
| A549 | RDK | RandomForestClassifier | 0.600 | 0.825 | 0.692 | 0.736 | 0.438 | 0.643 | 0.761 |
| H1.hESC | CDK | RandomForestClassifier | 0.500 | 0.785 | 0.607 | 0.671 | 0.297 | 0.548 | 0.708 |
| HeLa.S3 | PKNC | GaussianNB | 0.826 | 0.593 | 0.284 | 0.631 | 0.310 | 0.422 | 0.779 |
| **HepG2** | RDK | GaussianNB | 0.429 | 0.847 | 0.511 | 0.733 | 0.292 | 0.466 | 0.718 |
| **HT1080** | CDK | MLPClassifier | 0.517 | 0.769 | 0.633 | 0.659 | 0.296 | 0.569 | 0.728 |
| **HUVEC** | PKNC | SVC | 0.032 | 0.964 | 0.250 | 0.706 | -0.011 | 0.056 | 0.756 |
| **MCF.7** | CDK | GaussianNB | 0.452 | 0.809 | 0.437 | 0.721 | 0.259 | 0.444 | 0.724 |
| **NCLH460** | PKNC | SVC | 0.000 | 1.000 | 0.000 | 0.826 | 0.000 | 0.000 | 0.723 |
| **NHEK** | CDK | SVC | 0.000 | 1.000 | 0.000 | 0.742 | 0.000 | 0.000 | 0.643 |
| **SK.MEL.5** | CDK | XGBClassifier | 0.063 | 0.950 | 0.250 | 0.763 | 0.023 | 0.100 | 0.676 |
| **SK.N.DZ** | CDK | MLPClassifier | 0.545 | 0.692 | 0.529 | 0.635 | 0.237 | 0.537 | 0.679 |
| **SK.N.SH** | PDNC | QuadraticDiscriminantAnalysis | 0.547 | 0.732 | 0.422 | 0.683 | 0.259 | 0.476 | 0.703 |
| **GM12878** | PKNC | GradientBoostingClassifier | 0.133 | 0.939 | 0.400 | 0.752 | 0.115 | 0.200 | 0.687 |
| **K562** | ALL_COMP | DecisionTreeClassifier | 0.452 | 0.788 | 0.463 | 0.692 | 0.243 | 0.458 | 0.620 |
| **IMR.90** | PKNC | AdaBoostClassifier | 0.448 | 0.735 | 0.591 | 0.603 | 0.192 | 0.510 | 0.668 |
| **Average** |  |  | 0.370 | 0.829 | 0.405 | 0.704 | 0.197 | 0.362 | 0.705 |

**Table 5.** Performance of ML models on correlation features.

| **Cell-lines** | **Feature used** | **ML Model used** | **Sensitivity** | **Specificity** | **Precision** | **Accuracy** | **MCC** | **F1-score** | **AUC** |
| --- | --- | --- | --- | --- | --- | --- | --- | --- | --- |
| **A549** | PC_PDNC | GradientBoostingClassifier | 0.567 | 0.803 | 0.654 | 0.709 | 0.381 | 0.607 | 0.757 |
| **H1.hESC** | PDNC | RandomForestClassifier | 0.495 | 0.811 | 0.635 | 0.685 | 0.324 | 0.556 | 0.720 |
| **HeLa.S3** | PKNC | GaussianNB | 0.783 | 0.585 | 0.269 | 0.617 | 0.272 | 0.400 | 0.779 |
| **HepG2** | MAC | GaussianNB | 0.679 | 0.567 | 0.369 | 0.597 | 0.218 | 0.478 | 0.715 |
| **HT1080** | SC_PTNC | GaussianProcessClassifier | 0.583 | 0.769 | 0.660 | 0.688 | 0.359 | 0.619 | 0.738 |
| **HUVEC** | PDNC | AdaBoostClassifier | 0.302 | 0.897 | 0.528 | 0.732 | 0.243 | 0.384 | 0.757 |
| **MCF.7** | SC_PDNC | SVC | 0.000 | 1.000 | 0.000 | 0.754 | 0.000 | 0.000 | 0.753 |
| **NCL.H460** | PDNC | SVC | 0.000 | 1.000 | 0.000 | 0.826 | 0.000 | 0.000 | 0.683 |
| **NHEK** | PKNC | AdaBoostClassifier | 0.231 | 0.866 | 0.375 | 0.702 | 0.116 | 0.286 | 0.635 |
| **SK.MEL.5** | PDNC | GradientBoostingClassifier | 0.063 | 0.933 | 0.200 | 0.750 | -0.007 | 0.095 | 0.654 |
| **SK.N.DZ** | TAC | SVC | 0.424 | 0.865 | 0.667 | 0.694 | 0.327 | 0.519 | 0.739 |
| **SK.N.SH** | PC_PTNC | GaussianNB | 0.750 | 0.531 | 0.364 | 0.588 | 0.248 | 0.490 | 0.689 |
| **GM12878** | PC_PDNC | XGBClassifier | 0.350 | 0.899 | 0.512 | 0.771 | 0.288 | 0.416 | 0.715 |
| **K562** | DACC | AdaBoostClassifier | 0.405 | 0.808 | 0.459 | 0.692 | 0.221 | 0.430 | 0.642 |
| **IMR.90** | PC_PTNC | AdaBoostClassifier | 0.621 | 0.588 | 0.563 | 0.603 | 0.208 | 0.590 | 0.655 |
| **Average** |  |  | 0.417 | 0.795 | 0.417 | 0.694 | 0.213 | 0.391 | 0.709 |

### Models based on embeddings from DNABERT-2

Embeddings from large language models are known to encapsulate not just the individual bases, but also their biological significance in terms of structure, function, and interactions. In this approach, we generated high level representations of lncRNA sequences using both the pre-trained as well as the fine-tuned models. These embeddings were used to train ML models and the models were evaluated on the validation dataset. In the case of pre-trained embeddings, the model achieved an average AUC of 0.6586 and an average MCC of 0.1182. When fine-tuned embeddings were used as features, the performance of the model improved marginally, achieving an average AUC of 0.6604 and an average MCC of 0.1740. Detailed results for the performance of ML models on the validation dataset using pre-trained as well as fine-tuned embeddings as features are provided in Table 6 and Table 7 respectively.

**Table 6.** Performance of ML models on embeddings from pre-trained DNABERT-2 model.

| Cell-lines | Feature used | Sensitivity | Specificity | Precision | Accuracy | MCC | F1-score | AUC |
| --- | --- | --- | --- | --- | --- | --- | --- | --- |
| **A549** | GaussianNB | 0.678 | 0.555 | 0.500 | 0.604 | 0.228 | 0.575 | 0.689 |
| **H1.hESC** | MLP Classifier | 0.588 | 0.638 | 0.519 | 0.618 | 0.223 | 0.552 | 0.663 |
| **HeLa.S3** | MLP Classifier | 0.000 | 1.000 | 0.000 | 0.837 | 0.000 | 0.000 | 0.704 |
| **HepG2** | MLP Classifier | 0.000 | 1.000 | 0.000 | 0.728 | 0.000 | 0.000 | 0.694 |
| **HT1080** | MLP Classifier | 0.383 | 0.885 | 0.719 | 0.667 | 0.315 | 0.500 | 0.751 |
| **HUVEC** | MLP Classifier | 0.270 | 0.879 | 0.459 | 0.711 | 0.180 | 0.340 | 0.700 |
| **MCF.7** | MLP Classifier | 0.071 | 0.969 | 0.429 | 0.748 | 0.088 | 0.122 | 0.671 |
| **NCLH460** | Quadratic Discriminant Analysis | 0.375 | 0.697 | 0.207 | 0.641 | 0.059 | 0.267 | 0.536 |
| **NHEK** | MLP Classifier | 0.026 | 0.955 | 0.167 | 0.715 | -0.043 | 0.044 | 0.589 |
| **SK.MEL.5** | KNN | 0.063 | 1.000 | 1.000 | 0.803 | 0.224 | 0.118 | 0.578 |
| **SK.N.DZ** | MLP Classifier | 0.333 | 0.673 | 0.393 | 0.541 | 0.007 | 0.361 | 0.644 |
| **SK.N.SH** | MLP Classifier | 0.016 | 0.994 | 0.500 | 0.737 | 0.049 | 0.030 | 0.642 |
| **GM12878** | MLP Classifier | 0.117 | 0.965 | 0.500 | 0.767 | 0.152 | 0.189 | 0.678 |
| **K562** | Random Forest Classifier | 0.000 | 1.000 | 0.000 | 0.712 | 0.000 | 0.000 | 0.650 |
| **IMR.90** | LR | 0.552 | 0.735 | 0.640 | 0.651 | 0.292 | 0.593 | 0.688 |
| **Average** |  | 0.231 | 0.863 | 0.402 | 0.699 | 0.118 | 0.246 | 0.659 |

**Table 7.** Performance of ML models on embeddings from fine-tuned DNABERT-2 model.

| Cell-lines | ML model used | Sensitivity | Specificity | Precision | Accuracy | MCC | F1-score | AUC |
| --- | --- | --- | --- | --- | --- | --- | --- | --- |
| **A549** | SVC_linear | 0.000 | 1.000 | 0.000 | 0.604 | 0.000 | 0.000 | 0.669 |
| **H1.hESC** | Gradient Boosting Classifier | 0.456 | 0.798 | 0.600 | 0.661 | 0.271 | 0.518 | 0.688 |
| **HeLa.S3** | MLP Classifier | 0.174 | 0.983 | 0.667 | 0.851 | 0.287 | 0.276 | 0.737 |
| **HepG2** | Gradient Boosting Classifier | 0.161 | 0.927 | 0.450 | 0.718 | 0.131 | 0.237 | 0.659 |
| **HT1080** | SVC_linear | 0.417 | 0.769 | 0.581 | 0.616 | 0.199 | 0.485 | 0.696 |
| **HUVEC** | MLP Classifier | 0.302 | 0.879 | 0.487 | 0.719 | 0.214 | 0.373 | 0.727 |
| **MCF.7** | GaussianNB | 0.393 | 0.837 | 0.440 | 0.727 | 0.239 | 0.415 | 0.698 |
| **NCLH460** | XGBoost Classifier | 0.000 | 0.934 | 0.000 | 0.772 | -0.110 | 0.000 | 0.590 |
| **NHEK** | XGBoost Classifier | 0.128 | 0.946 | 0.455 | 0.735 | 0.126 | 0.200 | 0.644 |
| **SK.MEL.5** | AdaBoost Classifier | 0.250 | 0.933 | 0.500 | 0.789 | 0.244 | 0.333 | 0.491 |
| **SK.N.DZ** | XGBoost Classifier | 0.515 | 0.788 | 0.607 | 0.682 | 0.315 | 0.557 | 0.727 |
| **SK.N.SH** | XGBoost Classifier | 0.156 | 0.955 | 0.556 | 0.745 | 0.188 | 0.244 | 0.632 |
| **GM12878** | LR | 0.217 | 0.934 | 0.500 | 0.767 | 0.212 | 0.302 | 0.692 |
| **K562** | XGBoost Classifier | 0.238 | 0.885 | 0.455 | 0.699 | 0.155 | 0.313 | 0.598 |
| **IMR.90** | GaussianNB | 0.552 | 0.588 | 0.533 | 0.571 | 0.140 | 0.542 | 0.656 |
| **Average** |  | 0.264 | 0.877 | 0.455 | 0.711 | 0.174 | 0.320 | 0.660 |

### Fine-tuned DNABERT-2 model

In this approach, we used our training dataset to fine-tune the model and generate a fine-tuned tokenizer and model. Using this tokenizer and model, we generate high level representations of our lncRNA sequences and these representations are used by the model to generate predictions. The fine-tuned DNABERT2 model could not be evaluated while training as we were not able to implement five-fold cross validation. The fine-tuned model was evaluated on the validation dataset and performance metrics for the same are reported in Table 8.

**Table 8.** Performance of fine-tuned DNABERT-2 model on the validation dataset.

| **Cell-lines** | **Sensitivity** | **Specificity** | **Precision** | **Accuracy** | **MCC** | **F1-score** | **AUC** |
| --- | --- | --- | --- | --- | --- | --- | --- |
| **A549** | 0.500 | 0.861 | 0.703 | 0.718 | 0.393 | 0.584 | 0.762 |
| **H1.hESC** | 0.520 | 0.713 | 0.546 | 0.636 | 0.235 | 0.533 | 0.644 |
| **HeLa.S3** | 0.000 | 1.000 | 0.000 | 0.837 | 0.000 | 0.000 | 0.527 |
| **HepG2** | 0.000 | 1.000 | 0.000 | 0.728 | 0.000 | 0.000 | 0.698 |
| **HT1080** | 0.567 | 0.731 | 0.618 | 0.659 | 0.301 | 0.591 | 0.676 |
| **HUVEC** | 0.159 | 0.976 | 0.714 | 0.750 | 0.251 | 0.260 | 0.710 |
| **MCF.7** | 0.000 | 1.000 | 0.000 | 0.754 | 0.000 | 0.000 | 0.562 |
| **NCL.H460** | 0.000 | 1.000 | 0.000 | 0.826 | 0.000 | 0.000 | 0.576 |
| **NHEK** | 0.000 | 1.000 | 0.000 | 0.742 | 0.000 | 0.000 | 0.564 |
| **SK.MEL.5** | 0.000 | 1.000 | 0.000 | 0.789 | 0.000 | 0.000 | 0.468 |
| **SK.N.DZ** | 0.515 | 0.885 | 0.739 | 0.741 | 0.439 | 0.607 | 0.791 |
| **SK.N.SH** | 0.000 | 1.000 | 0.000 | 0.737 | 0.000 | 0.000 | 0.589 |
| **GM12878** | 0.000 | 1.000 | 0.000 | 0.767 | 0.000 | 0.000 | 0.718 |
| **K562** | 0.190 | 0.885 | 0.400 | 0.685 | 0.099 | 0.258 | 0.570 |
| **IMR.90** | 0.655 | 0.529 | 0.543 | 0.587 | 0.185 | 0.594 | 0.649 |
| **Average** | 0.207 | 0.905 | 0.284 | 0.730 | 0.127 | 0.228 | 0.634 |

### Cross-cell line prediction and fine-tuning limitations in DNABERT-2 for lncRNA classification

We expect the models trained on a specific cell line to perform well when applied to their own type. However, we wanted to test the theory put forward by Lin et al. [33630066], where they proposed that each cell type should be trained separately and that cross-cell type prediction would not be as accurate. To verify this, we evaluated a model (trained on the A549 cell line) against 14 other human cell lines, whose data was available in the lncAtlas database. The performance metrics for this approach are reported in Table 9. It was observed that our fine-tuned DNABERT-2 model performs best against A549, reporting an AUC of 0.7619, but it also performs significantly well (AUC > 0.7) for four other cell lines (i.e., HepG2, HUVEC, SK.N.DZ, GM12878). This indicates that the model has decent cross-cell line accuracy for some cell lines. However, to achieve optimal performance, each cell line must be trained separately and evaluated on its own validation dataset. To achieve this, we further fine-tuned the DNABERT-2 model that was already fine-tuned on the A549 cell line using each cell line’s individual training dataset. The performance metrics for this training approach are provided in Table 10. However, in this case, we observed that the average AUC decreased, suggesting that the DNABERT-2 model is not able to adequately capture the unique patterns in lncRNA specific to each cell line. This result indicates that while cross-cell line predictions can be moderately successful, fine-tuning models for individual cell lines may introduce overfitting or fail to generalize well due to limited data specific to each cell line.

**Table 9.** Performance of fine-tuned DNABERT-2 model (trained on A549 cell-line) evaluated on the validation dataset of other cell-lines.

| **Cell-lines** | **Sensitivity** | **Specificity** | **Precision** | **Accuracy** | **MCC** | **F1-score** | **AUC** |
| --- | --- | --- | --- | --- | --- | --- | --- |
| **A549** | 0.500 | 0.861 | 0.703 | 0.718 | 0.393 | 0.584 | 0.762 |
| **H1.hESC** | 0.328 | 0.860 | 0.609 | 0.648 | 0.224 | 0.427 | 0.668 |
| **HeLa.S3** | 0.391 | 0.695 | 0.200 | 0.645 | 0.068 | 0.265 | 0.553 |
| **HepG2** | 0.518 | 0.813 | 0.509 | 0.733 | 0.329 | 0.513 | 0.762 |
| **HT1080** | 0.417 | 0.833 | 0.658 | 0.652 | 0.277 | 0.510 | 0.654 |
| **HUVEC** | 0.492 | 0.855 | 0.564 | 0.754 | 0.362 | 0.525 | 0.751 |
| **MCF.7** | 0.440 | 0.794 | 0.411 | 0.707 | 0.229 | 0.425 | 0.670 |
| **NCI.H460** | 0.125 | 0.763 | 0.100 | 0.652 | -0.103 | 0.111 | 0.490 |
| **NHEK** | 0.256 | 0.741 | 0.256 | 0.616 | -0.003 | 0.256 | 0.561 |
| **SK.MEL.5** | 0.313 | 0.733 | 0.238 | 0.645 | 0.042 | 0.270 | 0.591 |
| **SK.N.DZ** | 0.455 | 0.846 | 0.652 | 0.694 | 0.330 | 0.536 | 0.754 |
| **SK.N.SH** | 0.328 | 0.754 | 0.323 | 0.642 | 0.082 | 0.326 | 0.612 |
| **GM12878** | 0.467 | 0.788 | 0.400 | 0.713 | 0.242 | 0.431 | 0.700 |
| **K562** | 0.429 | 0.740 | 0.400 | 0.651 | 0.166 | 0.414 | 0.617 |
| **IMR.90** | 0.379 | 0.647 | 0.478 | 0.524 | 0.027 | 0.423 | 0.650 |
| **Average** | 0.389 | 0.782 | 0.433 | 0.666 | 0.178 | 0.401 | 0.653 |

**Table 10.** Performance of fine-tuned DNABERT-2 model (initially trained on A549 cell-line, then trained on individual cell-lines) evaluated on their own validation dataset.

| **Cell-lines** | **Sensitivity** | **Specificity** | **Precision** | **Accuracy** | **MCC** | **F1-score** | **AUC** |
| --- | --- | --- | --- | --- | --- | --- | --- |
| **A549** | 0.556 | 0.774 | 0.617 | 0.687 | 0.336 | 0.585 | 0.734 |
| **H1.hESC** | 0.078 | 0.964 | 0.593 | 0.611 | 0.093 | 0.139 | 0.512 |
| **HeLa.S3** | 0.000 | 0.992 | 0.000 | 0.830 | -0.037 | 0.000 | 0.676 |
| **HepG2** | 0.161 | 0.960 | 0.600 | 0.743 | 0.207 | 0.254 | 0.741 |
| **HT1080** | 0.733 | 0.526 | 0.543 | 0.616 | 0.261 | 0.624 | 0.642 |
| **HUVEC** | 0.317 | 0.867 | 0.476 | 0.715 | 0.212 | 0.381 | 0.748 |
| **MCF.7** | 0.310 | 0.895 | 0.491 | 0.751 | 0.243 | 0.380 | 0.688 |
| **NCI.H460** | 0.000 | 1.000 | 0.000 | 0.826 | 0.000 | 0.000 | 0.525 |
| **NHEK** | 0.128 | 0.920 | 0.357 | 0.715 | 0.072 | 0.189 | 0.595 |
| **SK.MEL.5** | 0.000 | 1.000 | 0.000 | 0.789 | 0.000 | 0.000 | 0.531 |
| **SK.N.DZ** | 0.485 | 0.712 | 0.516 | 0.624 | 0.199 | 0.500 | 0.723 |
| **SK.N.SH** | 0.000 | 1.000 | 0.000 | 0.737 | 0.000 | 0.000 | 0.596 |
| **GM12878** | 0.033 | 0.995 | 0.667 | 0.771 | 0.111 | 0.063 | 0.682 |
| **K562** | 0.310 | 0.894 | 0.542 | 0.726 | 0.249 | 0.394 | 0.603 |
| **IMR.90** | 0.483 | 0.676 | 0.560 | 0.587 | 0.162 | 0.519 | 0.628 |
| **Average** | 0.240 | 0.878 | 0.397 | 0.715 | 0.141 | 0.268 | 0.642 |

### Performance comparison of CytoLNCpred and existing state-of-the-art classifiers

To further illustrate the efficacy of our method, we conduct a comparative analysis with other cutting-edge classifiers. Specifically, we evaluate existing predictors, namely lncLocator 2.0 and TACOS, which employ predictive algorithms to predict subcellular location of lncRNAs in different cell-lines.

Among these predictors, lncLocator 2.0 relies on the word embeddings and a MultiLayer Perceptron Regressor to predict CNRCI values. The predicted CNRCI values were then converted to labels using a fixed threshold value. The second predictor, TACOS, generated a variety of feature encodings using composition and physicochemical properties and tree-based algorithms were deployed to make the predictions. It is important to note that TACOS has been trained on 10 out of the 15 cell-lines. For a fair performance comparison, we leverage the performance metrics of evaluated on the validation dataset. Table 11 summarizes the evaluation of CytoLNCpred and other existing tools based on AUROC.

**Table 11.** Comparing the performance of our method and other existing classifiers using our validation dataset based on AUROC for all cell-lines.

|  | **IncLocator**<br>**2.0** | **TACOS** | **CytoLncPred –**<br>**Composition**<br>**features** | **CytoLncPred –**<br>**Correlation**<br>**features** |
| --- | --- | --- | --- | --- |
| **A549** | 0.592 | 0.741 | 0.761 | 0.757 |
| **H1.hESC** | 0.649 | 0.752 | 0.708 | 0.720 |
| **HeLa.S3** | 0.492 | 0.721 | 0.779 | 0.779 |
| **HepG2** | 0.487 | 0.712 | 0.718 | 0.715 |
| **HT1080** | 0.597 | 0.727 | 0.728 | 0.738 |
| **HUVEC** | 0.500 | 0.721 | 0.756 | 0.757 |
| **MCF.7** | 0.530 | - | 0.724 | 0.753 |
| **NCI.H460** | 0.500 | - | 0.723 | 0.683 |
| **NHEK** | 0.519 | 0.623 | 0.643 | 0.635 |
| **SK.MEL.5** | 0.500 | 0.567 | 0.676 | 0.654 |
| **SK.N.DZ** | 0.500 | - | 0.679 | 0.739 |
| **SK.N.SH** | 0.508 | 0.633 | 0.703 | 0.689 |
| **GM12878** | 0.500 | 0.568 | 0.687 | 0.715 |
| **K562** | 0.500 | - | 0.620 | 0.642 |
| **IMR.90** | 0.471 | - | 0.668 | 0.655 |
| **Average** | 0.523 | 0.676 | 0.705 | 0.709 |

## DISCUSSION

In recent years, researchers have recognized that the subcellular localization of lncRNAs plays a pivotal role in understanding their function. Unlike protein-coding genes, lncRNAs do not encode proteins directly. Instead, they exert their effects through diverse mechanisms, including interactions with chromatin, RNA molecules, and proteins. The precise localization of lncRNAs within the cell provides crucial information about their regulatory roles.

Subcellular localization of lncRNA gains prominence in recent times due to their role in gene regulation within the cell. A large number of aptamer and ASO based drugs are being developed using RNA nanotechnology. In recent years, the convergence of nanotechnology and long non-coding RNAs (lncRNAs) has yielded exciting developments in drug development. Nanoparticles, such as liposomes and exosomes, are being harnessed for targeted delivery of lncRNA-based therapeutics to cancer cells. Additionally, CRISPR-Cas9 technology, delivered via nanoparticles, enables precise gene editing by modulating lncRNA expression. Computational models and deep learning approaches are aiding our understanding of lncRNA-mediated mechanisms. Overall, this interdisciplinary field holds immense promise for personalized medicine, improved therapies, and better patient outcomes.

In recent times large language models are considered as SOTA methods and apart from the classical composition and correlation feature-based model, we also implemented DNABERT-2 for our classification problem. The DNABERT-2 model has been trained on the genomes of a wide variety of species and is computationally very efficient. DNABERT-2 uses Byte Pair Encoding to generate tokens which is known to perform better than k-mer tokenization. So, in order to fully exploit the DNABERT-2 model, we generated embeddings from both pre-trained and fine-tuned models. These embeddings when combined with ML methods were able to predict subcellular localization very well but poorer than a fine-tuned DNABERT-2 model. In our study, we compared the performance of DNABERT-2 with traditional composition and correlation-based features for classifying subcellular localization of lncRNAs. While DNABERT-2, a pre-trained language model, showed promising results, we found that traditional machine learning models trained on carefully crafted composition and correlation features consistently outperformed DNABERT-2. This suggests that for this specific task, the carefully engineered features capture the relevant biological information more effectively than the general-purpose representations learned by DNABERT-2.

Despite its promising potential, DNABERT-2 faces several challenges that limit its applicability for binary prediction. One of the most significant issues is the model’s lack of interpretability. The high-dimensional embeddings generated by DNABERT-2 can make it difficult to understand the rationale behind its predictions, hindering our ability to trust and utilize its outputs effectively. Moreover, the computational demands of DNABERT-2, especially when dealing with large nucleotide sequences, can be prohibitive for many applications. Additionally, the model’s reliance on extensive, annotated datasets can be a constraint, particularly for less well-studied organisms or specific biological phenomena.

A further limitation of DNABERT-2 is the variability observed in its training outcomes. This stochastic nature of LLM training can lead to different results even when trained on identical datasets and hyperparameters, hindering reproducibility and making it challenging to draw definitive conclusions from the model’s predictions. While techniques like setting random seeds and using fixed hyperparameters can mitigate this issue to some extent, it remains a fundamental challenge in LLM research and application. The complex interplay between the model’s architecture, training data, and hyperparameters can further contribute to the variability in outcomes, making it difficult to pinpoint the exact causes of discrepancies.

To address these limitations, future research should focus on developing techniques to improve the interpretability of DNABERT-2’s predictions. This could involve methods such as attention visualization or feature importance analysis. Furthermore, expanding the diversity of training data is essential to enhance the model’s generalizability across different biological contexts. By incorporating data from a wider range of organisms and conditions, DNABERT-2 could become a more versatile and reliable tool for genomic analysis. However, in our case, correlation-based features with classical machine learning algorithms outperformed all other approaches.

## CONCLUSION

Understanding the subcellular localization of lncRNA can provide great insights into their function within the cell. Computational tools have recently expanded the domain of subcellular localization by the development of faster and more accurate methods. In this study, we used a variety of machine learning as well as large language models to accurately predict lncRNA subcellular localization. The implementation of large language models to tackle biological problems is gaining momentum and our study also highlights its importance. The final model used in CytoLNCpred was designed using a traditional machine learning model trained using correlation-based features. This tool will help researchers to improve the functional annotation of lncRNA and use this information to develop RNA-based therapeutics.

## ACKNOWLEDGEMENTS

Authors are thankful to the Department of Science and Technology (DST-INSPIRE) and Indraprastha Institute of Information Technology, New Delhi, for fellowships and financial support. Authors are also thankful to Department of Computational Biology, IIITD New Delhi for infrastructure and facilities.

## AUTHORS’ CONTRIBUTIONS

SC collected and processed the datasets. SC implemented the algorithms and developed the prediction models. SC and GPSR analysed the results. SC created the back end of the web server, and SC and NKM created the front-end user interface. SC, NKM and GPSR penned the manuscript. GPSR conceived and coordinated the project. All authors have read and approved the final manuscript.

